# Choline monooxygenase transcript expression triggers glycine betaine accumulation in *Suaeda maritima* when subjected to a salt concentration optimal for its growth

**DOI:** 10.1101/314419

**Authors:** Sindhu Kuttan, H.M. Sankararamasubramanian, Ajay Kumar Parida

**Affiliations:** MS Swaminathan Research Foundation, Third cross road, Taramani Institutional Area, CPT Campus, Taramani, Chennai-600113, India

**Keywords:** Glycine betaine, HPTLC, NaCl, RNA expression, *Suaeda maritima*

## Abstract

*Suaeda maritima*(L.) Dumort, an annual halophyte known to be a salt-accumulator is also known for the accumulation of the osmolyte, glycine betaine (GB). This study is an attempt to understand the growth and GB accumulation under optimal concentration of 200 mM NaCl in *S. maritima.* Salt treatment with 200 mM NaCl showed a significant increase in shoot growth after two weeks. The shoots appeared succulent and turgid after two weeks of salt treatment compared to that of the control which appeared slender and wilted. The treated seedlings also exhibited a significant increase in GB content after two weeks of salt treatment. In order to determine the molecular basis of GB accumulation, qRT PCR of three key genes involved in the pathway, choline monooxygenase, betaine aldehyde dehydrogenase and phospho-ethanolamine N-methyl transferase was performed. Transcript level expression of the three genes revealed a high up-regulation of choline monooxygeanse transcripts when compared to that of the other two transcripts at two days of salt treatment. The results indicate that *S. maritima* requires salt for its growth and is a natural accumulator of GB. Although all the three genes were salt inducible, the high up-regulation of choline monooxygenase greatly contributes to the accumulation of GB under optimal growth as well as NaCl concentration.

**Abbreviations:**

BADH: betaine aldehyde dehydrogenase
CMO: choline monooxygenase
GB: glycine betaine
PEAMT: phosphoethanolamine N-methyl transferase

## Introduction

One of the major environmental factors that influence land ecosystems and consequently, agricultural productivity is soil salinity. Plants learn to cope with salinity in the soil by way of physiological and/or morphological adaptations, a striking example being the study of mangroves. Mangroves are morphologically and physiologically adapted to extreme environments as they experience conditions of high salinity, extreme tides, strong winds, high temperatures and muddy anaerobic soils and are able to take up water despite strong osmotic potentials in the inter-tidal regions (Kathiresan and Bingham, 2001). Maintenance of water balance under these conditions could be achieved only if tissue level water potential is lower than that of the substrate, and this is made possible in these plants by the accumulation of non-ionic, compatible solutes whose levels increase in the tissue with increasing salinity in the surrounding environment (Ball, 1988).

Glycine betaine (GB) is a commonly occurring betaine that confers osmotic tolerance in most plant species, particularly members of the spinach family (Adrian-Romero *et al.*, 1998). GB biosynthesis has been studied in different systems that accumulate it including bacteria, algae and higher plants. In plants, it is known to increase the osmolarity of cells as well as protect the structure of proteins involved in photosynthesis (Papageorgiou and Murata, 1995). Although the first step in GB biosynthetic pathway in plants involves the conversion of choline to betaine aldehyde by choline monooxygenase (CMO), which is followed by the conversion of betaine aldehyde to GB by Betaine aldehyde dehydrogenase (BADH), studies show that the conversion of ethanolamine to choline by the salt inducible Phospho ethanolamine N-methyl transferase (PEAMT) is crucial for the accumulation of GB, at least in plants that do not accumulate GB naturally (Mc Neil *et al,* 2001). CMO and BADH genes have been studied extensively in the case of spinach (Weretilnyk and Hanson, 1989; Burnet et al, 1995; Rathinasabapathi *et al*, 1997). The mangrove associates, *Suaeda* spp., and *Salicornia* spp., which belong to Chenopodiaceae, are known accumulators of GB (Hall *et al.,* 1978; Moghaieb *et al.,* 2004). *Suaeda maritima* (L.) Dumort, a halophyte growing in the estuarine regions is known to be a salt-accumulator, and grows well at a concentration of 200 mM NaCl (Clipson, 1987). Physiological and anatomical features of *Suaeda maritima* with respect to salinity tolerance have been studied extensively (Harvey et al, 1976; Hajibagheri et al, 1985; Flowers and Dalmond, 1992).

The present study is a preliminary attempt to study the genes involved in glycine betaine pathway in order to understand the mechanisms involved in the accumulation of GB in *Suaeda maritima*. An attempt has been made in this study to record the morphological changes with respect to salt treatment and also to find whether salt accumulation affects the physiological levels of GB in the plant. This was studied by analyzing the changes in transcript levels as well as by estimating the GB content, both at a salt concentration that is considered optimal for the growth of *S. maritima* (Clipson, 1987).

## Materials and methods

### Plant material and salt treatment

Seeds of *Suaeda maritima* were collected from Pichavaram mangroves, Tamil nadu, India, during late November. Seeds were germinated in pots containing vermiculite under distilled water. After one week of germination the seedlings were grown in modified Hoagland’s solution (Wang et al, 2007) inside a greenhouse under saturating daylight conditions (~925 μmol s^−1^ m^−2^, 30°C). One month old seedlings were transferred to hydroponics in 1X modified Hoagland’s solution and acclimatized for a week. The seedlings were then treated with 200 mM NaCl in modified Hoagland’s solution for 2 days, 4 days and 2 weeks. The concentration as well as duration of NaCl treatment was decided based on physiological changes reported in *S. maritima* under salt treatment by Clipson (1987). Seedlings maintained in modified Hoagland’s solution without NaCl served as control.

### Measurement of growth

For growth measurement analyses, fresh weight and dry weight measurements were performed using whole shoots of control and treated seedlings at 2 days and 2 weeks. After each treatment period, whole shoots were excised and weighed immediately. For dry weight measurement, the excised shoots were dried at 70°C for 48 hours and stored desiccated until weighing. The measurements were done for three biological replicates.

### GB extraction and purification

GB was extracted and purified according to Grieve and Grattan, (1983) and Bessiers *et al.,* (1999) respectively. About 200 mg dried, powdered plant material was suspended in 5ml de-ionized water and shaken overnight at 25°C, centrifuged at 300 g for 10 minutes at 4°C. The pellet was further re-suspended in 5 ml deionized water and extracted for another 24 hours as described above. The samples were then pooled, lyophilized and stored at−20°C until further use.

AG 1 X-8 resin (Bio Rad), Cl^−^ form, was converted to OH^−^ form by following the procedure described by the manufacturer. ~2.0ml of the resin was then packed in a 2ml syringe, which was dried down by centrifugation at 300 x g for 2 minutes at 4°C. 250μl of freeze-dried extract was loaded followed by the addition of 750μl de-ionized water. The samples were centrifuged at 300 x g for 3 minutes to collect the eluant containing GB and were used for HPTLC (High Performance Thin Layer Chromatography) analyses.

### GB estimation using HPTLC

#### GB standard preparation

Separation and detection of GB on TLC chromatoplates was performed according to Tyihák *et al.,* (1994) except that a HPTLC method was used instead of an OPLC (Over-pressured Layer Chromatography) method used by the authors. One hundred mg GB (Sigma, USA) was weighed and dissolved in 10ml de-ionized water. The sample was diluted to give 1mg/ml GB and spotted onto a 20cm x 10 cm TLC chromatoplate precoated with silica gel 60 F_254_ (Merck). Standard solutions corresponding to 2μg, 4μg, 6μg, 8μg, 10μg and 12μg GB was loaded onto the TLC plate using Linomat 5.0 sample applicator (CAMAG, Switzerland). The plate was then run in a TLC tank with a solvent mixture of n-propanol/methanol/0.1M Sodium acetate (2/3/3, v/v/v). Post chromatographic derivatization was performed after the plates were removed from the chamber after the solvent front reached up to approximately 0.5 to 1.0cm below the distal end of the plate. The plate was dried under cool air for 15 minutes, dipped in Dragendorff’s reagent (see below) for 3 seconds, and dried under a stream of air in a laminar air flow for 45 minutes. The spots developed were scanned in HPTLC densitometric scanner (CAMAG, Switzerland) using *winCATS planar chromatography manager* software (Fig. S1). The samples used for GB estimation were loaded in triplicates along with the standard. GB content was expressed in μmol per gram dry weight.

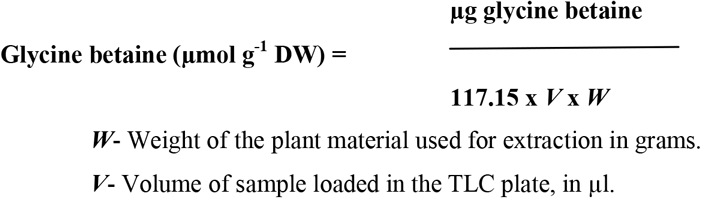

### Total RNA isolation and qRT PCR analyses

Total RNA was isolated from whole shoots of *S. maritima* using Trizol reagent (cat. #T9424, Sigma Chemical Co. USA), and the procedure was followed as described by the manufacturer. The isolated total RNA was subjected to DNAse (NEB) treatment and was further purified using RNA clean up kit (Macherey-Nagel, Germany). Purified RNA samples were then quantified using a UV-1601 spectrophotometer (Shimadzu, Japan) and the integrity of RNA samples were checked using agarose gel electrophortesis. RNA samples used for cDNA synthesis exhibited a 260/280 ratio between 1.8 to 2.0 and a 260/230 ratio greater than 1.7. cDNA synthesis (High Capacity cDNA Reverse Transcription kit, Applied Biosystems) was performed using 2μg each of the total RNA as the template. For qRT PCR analyses, 1.0μl each of the cDNAs was used as the template. *Actin* was used as the endogenous control. Primer sequences used for the amplification of *Actin, PEAMT* and *BADH* were synthesized as described by Sahu and Shaw (2009). Primers for *CMO* were designed from the *SmCMO* cDNA sequence (NCBI accession # JX629239) using Primer express software, version 3.0 (Applied Biosystems). *SmCMOfwd*: 5’- TGTTTTATTGCTTGTGTTCTTGGAA−3’; *SmCMOrev*: 5’GGGTGCTATGGTGGTGTTTATTTAT−3’. The cDNAs were normalized with *SmActin* primers and the amount of cDNA giving the same C_t_ value across the samples was used for bulk reaction. SYBR Green qRT-PCR assays were carried out using the Step One Plus^TM^ Real Time PCR system, according to the manufacturer’s instructions (Applied Biosystems). The reaction comprised 10 μl of Power SYBR Green PCR master mix (2X), 2.5 μl of 2μM forward and reverse primer mix to give a final concentration of 250 nM, 1 μl of cDNA from each sample and 17 μl of deionized H_2_O. The PCR cycling conditions were as follows: 95 °C for 10 min., followed by 40 cycles of 95 °C for 15 seconds and 60 °C for 1 min., in a 96-well optical reaction plate (Applied Biosystems). Assays were carried out in triplicates in order to evaluate data reproducibility. Specificity of the amplicons was verified by melt-curve analysis (60° to 95 °C) after 40 cycles. Data from three replicate qRT-PCR samples were analyzed using StepOne^TM^ software by comparative C_T_ (ΔΔC_t_) quantitation method. The ΔC_t_ (differences in the C_t_ value between the target and internal control) for each sample was subtracted from that of the calibrator (ΔΔC_t_). *SmCMO, SmBADH* and *SmPEAMT* expression levels were calculated using 2^−ΔΔCt^ and the values were represented as n-fold difference relative to the control. Each experiment was repeated three times with independent set of samples.

### Statistical analyses

All the data were calculated as mean ± standard error of the values obtained in all the above experiments. Statistical significance analyses used in the study was calculated using Student’s t-test. Values of *p* < 0.05 were considered significant.

## Results and Discussion

### Salt concentration and growth

*S. maritima* seedlings were subjected to salt treatment to record the morphological changes and to determine the influence of NaCl on their growth after two days and two weeks of treatment. The seedlings treated with 200 mM NaCl showed emergence of more number of new leaves compared to the control (Fig. 1d) by the end of 2 weeks. The leaves appeared more succulent compared to the control seedlings. The leaves of control seedlings were thinner, shorter and curled compared to the salt treated seedlings at the end of 2 weeks. The seedlings treated with NaCl for 2 days showed 39 and 42 percent reduction in fresh and dry weight respectively, whereas those treated with NaCl for 2 weeks showed a 69 and 56 percent increase in fresh and dry weight respectively compared to control seedlings (Table 1). The salt concentration used in the present study was 200 mM NaCl, chosen to assess the short term as well as the long-term effects of salinity in *S. maritima;* salt concentrations above 200 mM NaCl retard growth of the plant (Clipson, 1987). There is a significant increase in shoot growth at two weeks of salt treatment (Table 1), contradicting the results obtained by Moghaieb *et al.* (2004), who reported significant reduction in the growth of the seedlings beyond 85mM NaCl. Interestingly, a marked reduction in shoot growth was observed in the seedlings after two days of salt treatment (Table 1), which could be attributed to the recovery phase of the seedlings after a sudden increase in salt concentration. Clipson (1987) observed an increase in shoot growth only after 4 days of (200 mM) salt treatment.

**Fig 1:**
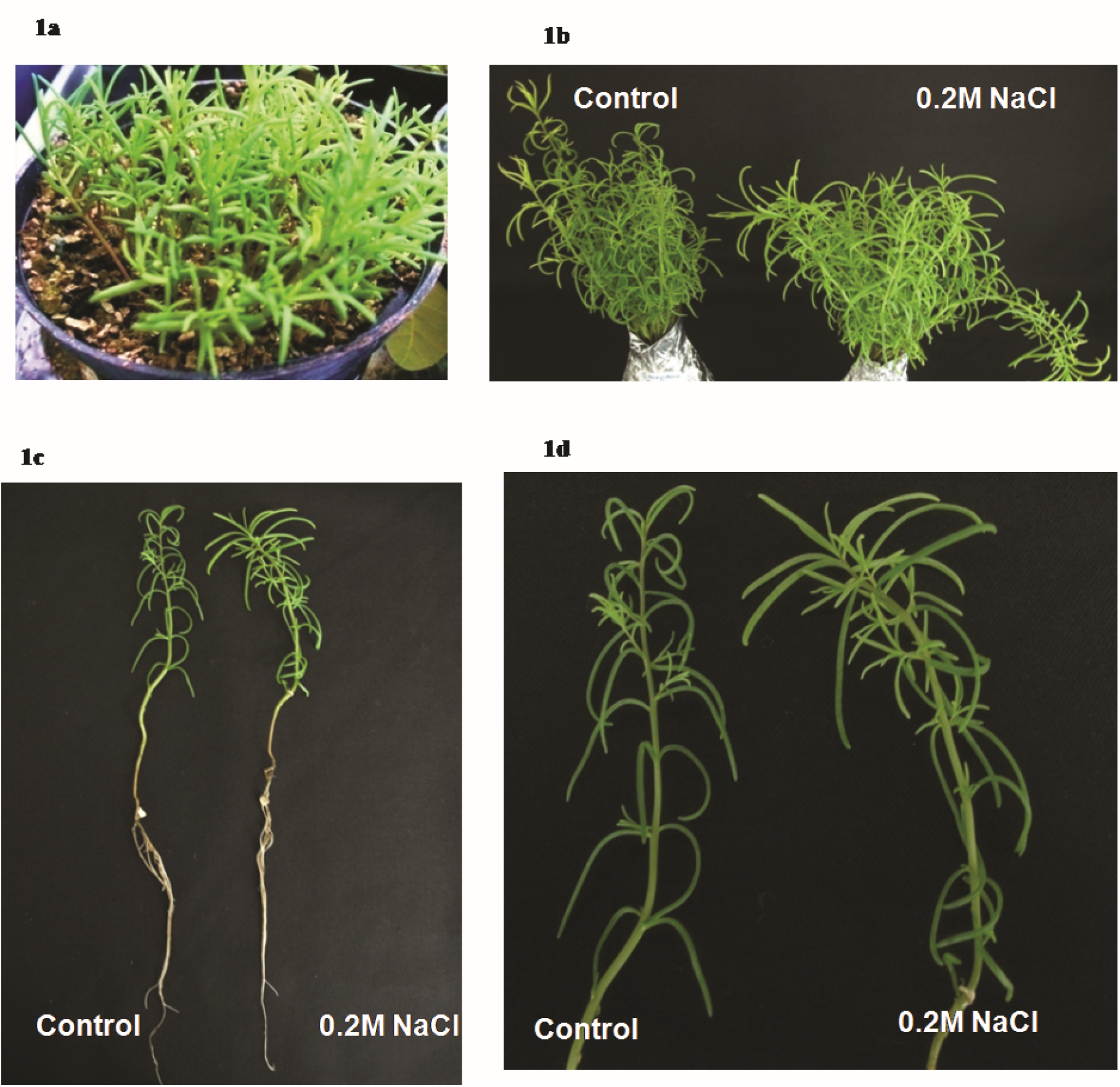
Salt treatment of *Suaeda maritima*. **(1a)** *Suaeda maritima* seeds germinated on vermiculite **(1b)** One month old *S.maritima* seedlings treated with 200mM NaCl **(1c)** Control and salt treated seedlings after 2 weeks of treatment **(1d)** Shoot region of control and salt treated seedlings after 2 weeks of treatment.

**Table 1.** Shoot growth measurements in *Suaeda maritma* seedlings treated with 200mM NaCl.

|  | (a) |  | (b) |  |
| --- | --- | --- | --- | --- |
|  | Whole shoot fresh weight |  | Whole shoot dry weight |  |
|  | g |  | g |  |
|  | 2 Days | 2 Weeks | 2 Days | 2 Weeks |
| **Control** | 5.2 ±0.2 | 6.6 ±0.4 | 0.8 ±0.3 | 1.1 ±0.05 |
| **NaCl treatment** | 3.1 ±0.3 | 11.1 ±0.3 | 0.35 ±0.03 | 1.7 ±0.03 |
| **P- value** | 0.0053 | 0.0007 | 0.2477 | 0.0033 |

### GB estimation by HPTLC

HPTLC (High Performance Thin Layer Chromatography) analysis of *S. maritima* purified samples showed the detection of peaks corresponding to that of the standard GB (Fig.S1a). TLC spots were also detected in the same R_f_ region as that of the standard (Fig. S1b,d). There was no significant difference between GB content of whole shoots of control and NaCl treated *S. maritima* (Table 2) after 2 days of salt treatment.

**Table 2.** Glycine betaine content in whole shoots of *Suaeda maritima*.

|  | Glycine betaine content<br>μmol/g dry wt. |  |
| --- | --- | --- |
|  | 2 Days | 2 Weeks |
| Control | 50.6 ± 4.7 | 50.7 ± 0.8 |
| NaCl Treatment | 48.7 ± 2.8 | 57.4 ± 1.1 |
| *P*- value | 0.9959 | 0.0045 |

GB content in control plants was 50.58 μmol per gram dry weight, whereas for the NaCl treated plants it was 48.65 μmol per gram dry weight. However, after 2 weeks of treatment, the GB content in NaCl treated plants was 15 percent more than that of untreated control plants (Table 2). GB content after two weeks of salt treatment was 57.4 μmol per gram dry weight; GB content in untreated control plants after two weeks of NaCl treatment was 50.7 μmol per gram dry weight. The extraction procedure of GB from dry plant material was followed according to Grieve and Grattan (1983). The anion exchange purification step (Bessieres *et al.,*1999), eliminates 80% of total amino acids, sugars etc. The purification step also eliminates proline, which accumulates in *S. maritima* in response to salt (Moghaieb *et al.,* 2004). By far, the most sensitive method to analyze GB content has been that by LC-MS where picogram levels of GB can be detected (Airs and Archer, 2010); the most economical being spectrophotometric detection (Grieve and Grattan, 1983; Stumpf, 1984). Other methods used for estimating betaine include NMR and HPLC (Bessiers *et al.*1999), GC (Hitz and Hanson, 1980) and FAB-MS (Rhodes *et al.,* 1987). GB was also quantified using OPLC (Over pressure layered chromatography), a method comparable to that of NMR (Tyihák *et al.,* 1994). This is the first report where HPTLC (High Performance Thin Layer Chromatography) is used for GB estimation (Fig. S1), the difference between this method and OPLC being the TLC plate run in a pressurized chamber in OPLC which is not the case in HPTLC. Nevertheless, it was possible to achieve comparable resolution of GB spots with HPTLC, and this could be attributed to the purification step. GB was the major, and visibly the sole spot detected in *S. maritima* after passing through the anion exchange resin (Fig. S1b).

### Transcript expression under salt treatment

qRT-PCR analysis showed that all the three transcripts, *SmCMO, SmBADH* and *SmPEAMT,* were upregulated by NaCl (200 mM) although basal levels of transcripts were found in the control plants (Fig. 2). *SmCMO* was significantly upregulated after 2 days of NaCl treatment when compared with that of *SmBADH* and *SmPEAMT,* whose transcript levels were almost the same after two days of NaCl treatment. *SmCMO* showed a 34-fold increase in the transcript levels when compared with the control plants after two days of NaCl treatment, whereas *SmPEAMT* and *SmBADH* showed 5-fold increase in transcript expression.

**Fig 2:**
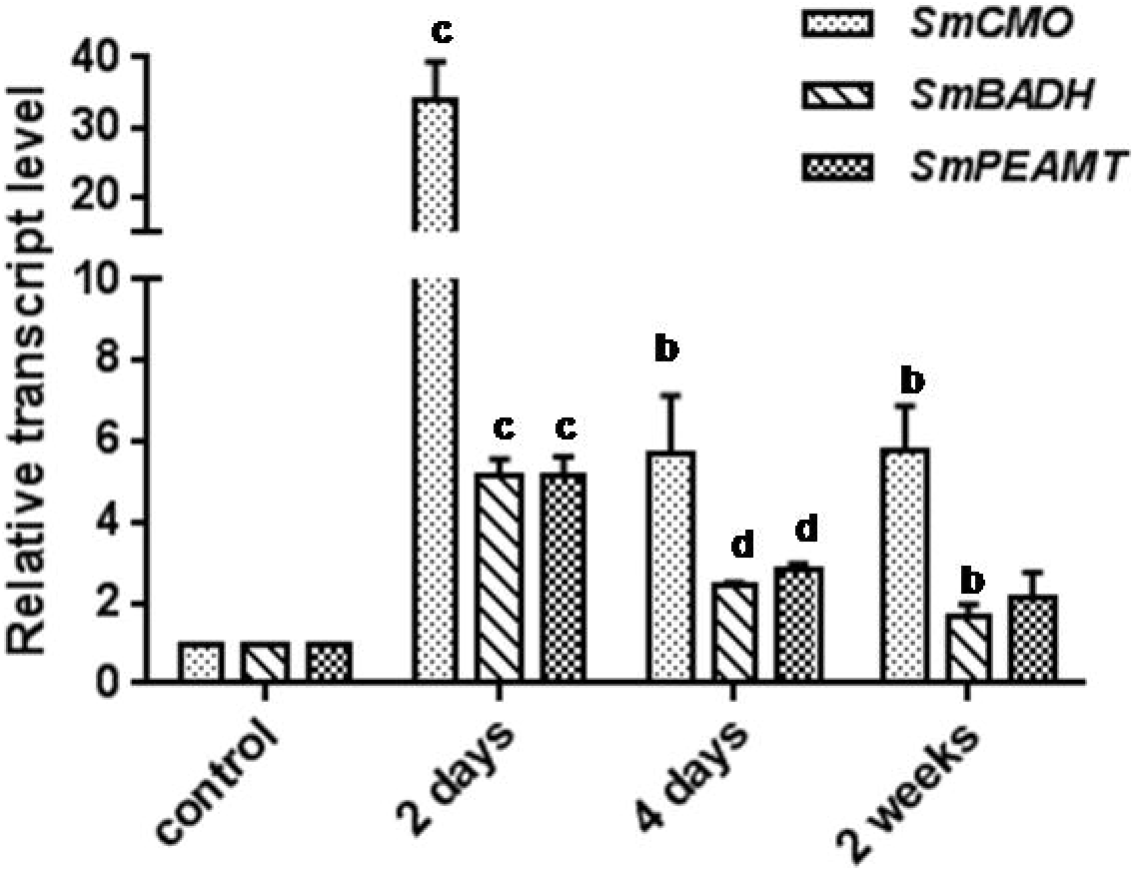
qRT-PCR of SmCMO, SmBADH and SmPEAMT. Whole shoots of one-month old S. maritima seedlings treated with 200mM NaCl were harvested at 2 days, 4 days and 2 weeks. qRT-PCR of the samples were performed with DNAse-free Total RNA samples. SmActin primers were used for endogenous control. The mean values between control and treated samples were tested using student’s t-test and the level of significance marked using the p-values: a, p ≤ 0.05 b, p ≤ 0.01 c, p ≤ 0.001 d, p ≤ 0.0001.

The high up-regulation of *SmCMO* transcripts under salt treatment of *S. maritima* (Fig. 2) is an indication that it triggers the pathway towards GB accumulation in response to salinity. In a study involving short term response of *S. maritima,* where the seedlings were treated with 425mM NaCl for 24 hours (Sahu and Shaw, 2009), the transcript levels of *SmPEAMT* were up-regulated by over 20-fold; there was no up-regulation in the *SmBADH* transcript levels. Interestingly, there was no mention of transcript levels of *SmCMO* in the study by the authors. High expression of *PEAMT* as well as *CMO* transcripts was also reported in *Atriplex nummularia* (Tabuchi et al., 2005). Perhaps, as indicated by Sahu and Shaw (2009), increase in choline levels is essential for the plants to accumulate enough GB and/or other osmolytes in the cytosol to counteract the effect of high concentrations of NaCl accumulated in the vacuole (Bohnert and Sheveleva, 1998). Moghaieb *et al.* (2004) have indicated in their study involving *S. maritima* that increasing levels of BADH transcripts were observed with increase in salt concentration, and that the results coincided with the increase in GB content. The marginal increase in GB at 2 weeks (Table 2) when the *SmCMO* transcript levels were still high compared to *SmBADH* and *SmPEAMT* (Fig.2) indicate that *SmCMO* and not SmPEAMT, plays a significant role in the accumulation of GB in *S. maritima.* The increase in *SmCMO* transcripts at 2 days, however, did not result in any increase in GB concentration (Fig.2, Table 2) upon salt treatment. The transcript level of *SmBADH* is not as high as that of *SmCMO* in 2 days or 2 weeks of salt treatment (Fig. 2). Although it has to be seen whether the pattern of transcript level expression of the genes would change under conditions above 200 mM NaCl, it appears that the increase in GB content is attributed to *SmCMO* rather than *SmBADH* (Fig.2).

## Conclusion

Based on the morphological observations and growth analyses, it could be said that *S. maritima* requires NaCl for optimum growth and to regain its morphology (leaf succulence) over time in the presence of optimal salt concentrations. There is a marginal increase in GB after two weeks of treatment, but also a considerable increase in shoot growth. This indicates that probably other osmolytes come in to play during salt treatment along with GB; S*. maritima* is also known to accumulate proline (Moghaieb *et al*., 2004). Although the marked increase in *SmCMO* transcript levels cannot be correlated to the GB content in the present study, it appears that the high up-regulation of *SmCMO* under salt conditions is essential for the resultant concentration of GB in the tissue. Further studies on the expression of *SmCMO* as well as GB content at increasing NaCl concentrations would throw more light on its expression in response to NaCl. It would be interesting to analyse the transcript expression levels of *SmBADH* and *SmPEAMT* at increasing NaCl concentrations.

## Acknowledgement

We thank Mr. Karthik and Mr. Muthu Kumar, Lichen ecology and bio-prospecting laboratory, MSSRF, Chennai, for their help in handling HPTLC instrument. We thank the Department of Biotechnology, India, for providing the funds to carry out the study.

## Competing interests

The authors declare no competing interests.

